# Epigenetic immune-modulation by Histone Deacetylase Activity (HDAC) of tissue and organ regeneration in *Xenopus laevis*

**DOI:** 10.1101/2020.02.05.936294

**Authors:** Nathalia Pentagna, Felipe Soares dos Santos, Fernanda Martins de Almeida, José Garcia Abreu, Michael Levin, Katia Carneiro

## Abstract

In the present work we propose to shed light on the epigenetic control of immune mechanisms involved during *Xenopus* tail regeneration. Here we show that the first 24 hour post amputation (hpa), which exclusively encompasses the first wave of myeloid differentiation, are crucial to epigenetically modulate the regenerative ability of *Xenopus* tadpoles. During this developmental window, HDAC activity was shown to be necessary for the proper establishment of myeloid cells dynamics in the regenerative bud, mainly contributing to modulate the behavior of monocytes/macrophages and neutrophils as well the mRNA expression pattern of the main myeloid markers, such as LURP, MPOX, Spib and mmp7. In addition, we functionally bridge the spatial and temporal dynamics of lipid droplets, the main platform of lipid mediators synthesis in myeloid cells during the inflammatory response, and the regenerative ability of *Xenopus* tadpoles showing that 15-LOX activity is a key player during tail regeneration. Taken together our results support a role for the epigenetic control of inflammatory response during tissue and organ regeneration, which may positively impact translational approaches for regenerative medicine.

**Summary statement:** We propose that Epigenetic mechanisms HDAC-dependent can control myeloid cells behavior upon tissue injury and that HDAC inhibitors may be used for tissue regeneration in translational studies.

## Introduction

Tissue and organ regeneration is a fascinating topic of study that has caught the attention of several generations of scientists. Different models of study have been used to unravel the cellular, genetic and molecular basis of animal regeneration. One of the most widely used model is the tail of *Xenopus laevis* tadpoles. In fact, between larval stages 40 and 44, swimming tadpoles have an amazing ability to regenerate tissues and organs upon tail amputation. About 72 hours post amputation, tadpoles regenerate notochord, muscles, epidermis and nervous system. Studying this regenerative capacity is extremely attractive as it resembles mammalian regeneration much more than the regenerative strategies in anurans, such as salamander, for example. For these reasons, this is a very tractable model to deep explore the main mechanisms involved in tissue and organ regeneration with critical implications for translational medicine.

Different studies have proposed that dormant developmental programs that tight control gene expression are re-activated during tissue and organ regeneration. In fact, it has been shown that key molecular pathways such as *TGF-β, BMP, Wnt*, (Beck, Christen, & Slack, 2003; Ho & Whitman, 2008; Sugiura, Tazaki, Ueno, Watanabe, & Mochii, 2009) along with bioelectrical signaling (Adams, Masi, & Levin, 2007; Beane, Morokuma, Lemire, & Levin, 2013; Levin, 2007, 2014; A. S. Tseng, Beane, Lemire, Masi, & Levin, 2010) are tight regulated upon *Xenopus* tail amputation, both in time and space, giving rise to a finely tuned program that drives full organ regeneration. Besides this remarkable feature, it has been proposed that cells from the immune system may also contribute to the regenerative potential of different organisms. For instance, cells from myeloid origin such as monocytes/machophages and neutrophils have been implicated in zebra fish, axolot and mouse regeneration (Aurora et al., 2014; Godwin, Pinto, & Rosenthal, 2013; Morales & Allende, 2019; Petrie et al., 2014).

Regardless myeloid cells have been described in *Xenopus*, evidences for their participation during tail regeneration are still lacking. As in mammals, *Xenopus* myeloid cells are first generated during embryonic stages during the so called first wave of differentiation, or primitive hematopoiesis, in a specialized tissue that is homolog to the mammalian yolk sac, the anterior ventral blood island (aVBI) at the neurula stage (Ciau-Uitz, Liu, & Patient, 2010; Ciau-Uitz, Walmsley, & Patient, 2000; Lane & Sheets, 2002; Lane & Smith, 1999). In addition to the primitive hematopoiesis, *Xenopus* also present a second wave of hematopoietic differentiation, the definitive hematopoiesis, that takes place in the dorso-lateral plate mesoderm (DLP) starting from stage 43 (Ciau-Uitz et al., 2000; Medvinsky & Dzierzak, 1996).

The master gene involved in myeloid differentiation is *Spib*, whose expression is first observed in the aVBI at stage 17 (Agricola et al., 2016; Costa, Soto, Chen, Zorn, & Amaya, 2008). Upon specification, *Spib+* cells start migrating away aiming to colonize the developing embryo. *Spib* is the *spi1/pu.1* zebra fish homolog and is necessary for hemangioblast-like cells differentiation into primitive myeloid precursor cells that will disperse into the embryo even before the raising of functional vessels (Costa et al., 2008). In fact, other myeloid markers such as mmp7, LURP and MPOX are downstream targets of *Spib* and contribute to different biological functions of myeloid cells, such as extracellular matrix remodeling, phagocytosis and immune defense (Harrison et al., 2004; Paredes, Ishibashi, Borrill, Robert, & Amaya, 2015; S. J. Smith, Kotecha, Towers, Latinkic, & Mohun, 2002). Costa et al have shown that *spib* positive cells are capable of migration towards an injury site during tailbud-stage, displaying in this way a functional behavior even in the absence of a functional vasculature. In this sense, macrophages and neutrophils are the two main myeloid population with relevant roles during regeneration.

In a previous work we have demonstrated the epigenetic control of *Xenopus* tail regeneration by Histone Deacetylase (HDAC) activity (A.-S. Tseng, Carneiro, Lemire, & Levin, 2011). In this work we showed that the treatment of regenerating swimming tadpoles, starting at stage 40, with HDAC inhibitors (iHDACs) impairs regeneration and leads to a lack of regenerative ability. In addition, we have recently shown that iHDAC treatment of mammalian myeloid cells both *in vitro* and *in vivo* disrupts myeloid differentiation favoring cell cycling in detriment of cell differentiation (Cabanel et al., 2015; Cabanel, da Costa, El-Cheikh, & Carneiro, 2019). As a consequence, myeloid progenitors display a high plastic phenotype favoring an anti-inflammatory functional profile. These findings are very interesting because, given the anti-inflammatory activity of HDAC inhibitors (iHDAC), we hypothesized that HDAC activity blockade might be dampening the inflammatory response from the onset, which in turn, would be required for regeneration. Thus, in the present paper, we hypothesized whether epigenetic mechanisms HDAC-dependent would emerge as putative candidates playing key roles as myeloid cells modulators for new strategies to improve regeneration in vertebrates. Therefore, iHDAC should be dampening the inflammatory response evoked by myeloid cells, which is also supposed to be necessary for *Xenopus* tail regeneration.

To test this hypothesis, we have focused on the characterization of myeloid sub-sets found in the regenerative bud upon tail amputation as well as on the characterization of their populational dynamics along the main regenerative events. Here we describe, for the first time, that myeloid cells belonging exclusively to the first wave of hematopoietic differentiation are the specific target of iHDACs. Upon tail amputation HDAC activity was necessary to proper modulate both myeloid cells behavior and the mRNA expression pattern of the myeloid markers mmp7, Spib, MPOX and LURP. Our results indicate that, similarly to mammalians, HDAC activity is a key player during myeloid differentiation with critical effects on the inflammatory response related to the regenerative ability during tissue and organ development. Our findings report for the first time the epigenetic control HDAC-dependent of the immune modulation of tissue and organ regenerative capacity with critical implications for regenerative medicine.

## Results

### HDAC activity is necessary for the regenerative ability exclusively during the first wave of myeloid differentiation

It has been shown that both VBI and DLP contribute with embryonic and definitive/adult blood cells, respectively. The VBI has been separated into two different territories: aVBI and pVBI, which give rise to erythrocytes and myeloid cells. pVBI also gives rise to short term lymphoid cells (T cells) (Costa et al., 2008; Rollins-Smith & Blair, 1990; P. B. Smith & Turpen, 1985; Tashiro, Sedohara, Asashima, Izutsu, & Maéno, 2006). DLP, in turn, gives rise to hematopoietic stem cells and therefore to all hematopoietic lineages in the adult (Chen & Turpen, 1995; Kau & Turpen, 1983). These two waves of hematopoietic differentiation take place at different times and at different sites into the embryo. While the first wave takes place at ventral sites of the embryo starting at stage 19, the second one starts only at stage 43 at dorsal sites. To better understand the epigenetic modulation of myeloid cells behavior during tail regeneration, we first investigated the requirement of HDAC activity during the first and/or the second waves of myeloid differentiation as well as its contribution to the regenerative ability. For this, tadpoles from different developmental stages, covering both the first or the second wave of myeloid differentiation, were incubated with iHDAC (Figure 1A). Interestingly incubation with iHDAC exclusively during the first 1, 3, 6, 12 or 24 hpa, starting at stage 40, was sufficient to gradually impair the regenerative ability observed in control group (Figure 1B, B’). In contrast, when tadpoles were incubated exclusively after the first 1, 3, 6, 12 and 24 hpa, until stage 47/48, the regenerative ability was gradually rescued (Figure 1C,C’).

**Figure 1:**
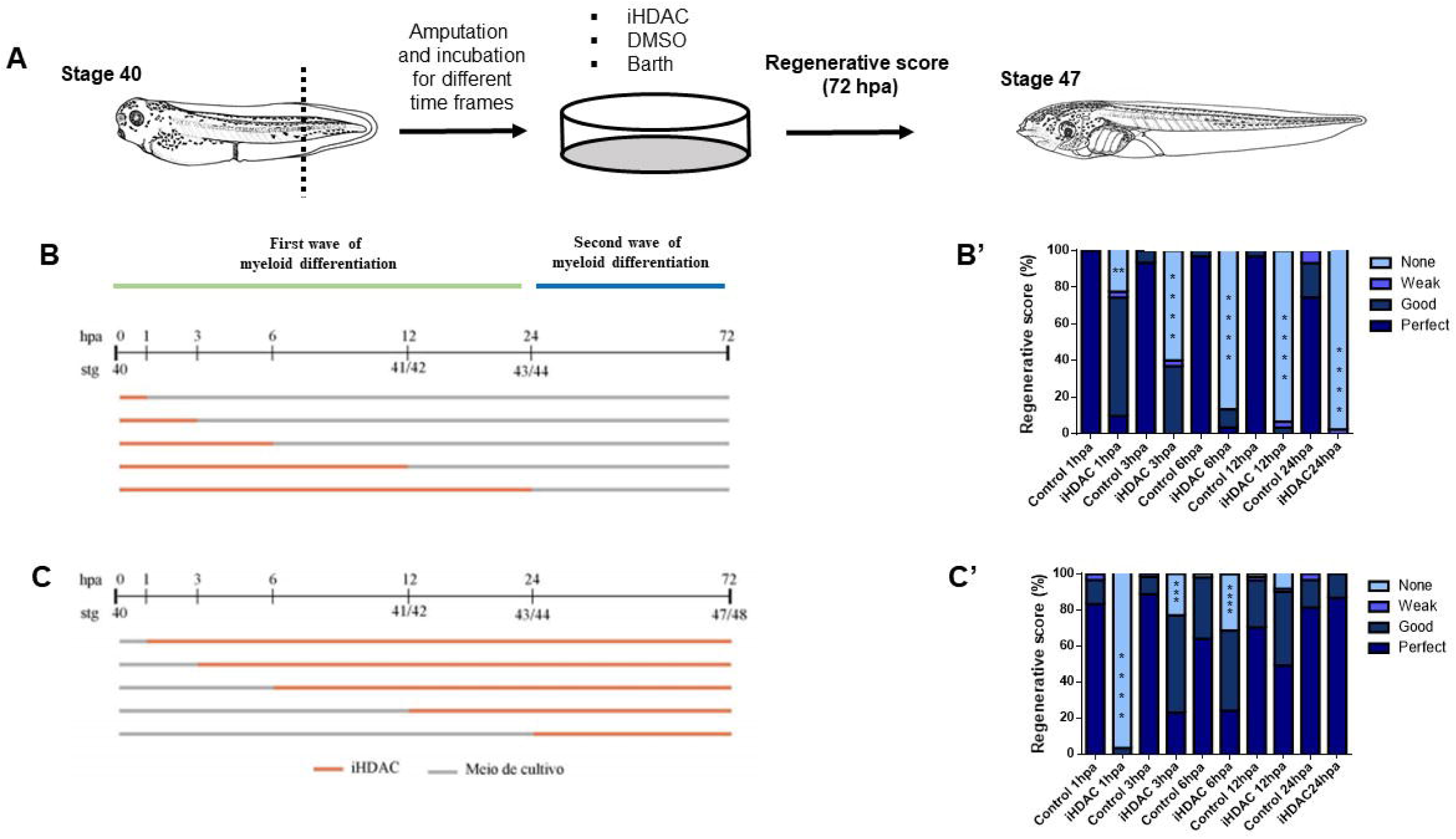
HDAC activity is required for tail regeneration exclusively within the first 24hpa that encompasses the first myeloid wave of differentiation. (A) At stage 40 tail was amputated and incubated with iHDAC during different windows of myeloid development (B,C). The tadpoles were aged untilstage 47/48 when were scored (B’,C’). For B the number of animals used in each treatment corresponds to 30, except for iHDAC 1h, with 31 animals and both groups 24hpa, with 43 animals. For C, the number of animals used in each treatment corresponds, respectively, to 60, 60, 62, 61, 50, 54, 57, 61, 59 and 60. Qui-square test was used. \*\*\**p*<0,001; \*\*\*\**p<*0,0001.

We conclude that the first 24 hpa, which exclusively encompasses the first wave of myeloid differentiation, are crucial to epigenetically modulate the regenerative ability of *Xenopus* tadpoles.

### The behavior of phagocytic cells positive for naphthyl esterase is modulated by HDAC activity at 24 hours post amputation

To gain insight into the epigenetic control of immune cells behavior upon tail amputation, we performed an analysis of immune cells dynamics into the regenerative bud by flow cytometry. We were able to determine 3 different cell populations that we named P1, P2 and P3 taking in account the parameters FSC (cell size) and SSC (cell complexity or relative granularity) (Figure 2). While P1 displays both low size and relative granularity and may correspond to lymphocytes, P3 characteristics may correspond to myeloid cells, most probably to granulocytes. Upon 24 hpa we observed that iHDAC treated tadpoles presented a consistent increase in the P3 sub-set mostly due to a decrease in P2 sub-set if compared to control group (Figure 2A,D,H,I). At 48 and 72 hpa, we did not observe any significant difference in cellular dynamics of P2 and P3 sub-sets (Figure 2B,C,E,F,H,I). In contrast, P1 sub-set dynamics was not disrupted along the time (Figure 2). Next, we asked whether HDAC activity is necessary for the global dynamics along the time frame of regeneration. For this, we compared cell dynamics upon tail amputation to “time zero/steady state” tails that were obtained from the tip of amputated tissues at the time of tail amputation. Interestingly, P2 sub-set increases significantly at 24 hpa and returns to the steady state already 48 hpa in control animals (Figure 2J). In contrast, P3 sub-set presented a complementary pattern in which at 24 hpa presented a significant decrease and return to steady state already at 48 hpa in control animals (Figure 2J). iHDAC treated tadpoles, on the other hand, presented a disruption in this pattern. In fact, P2 sub-set remained stable at 24 hpa instead of increasing (Figure 2J).

**Figure 2:**
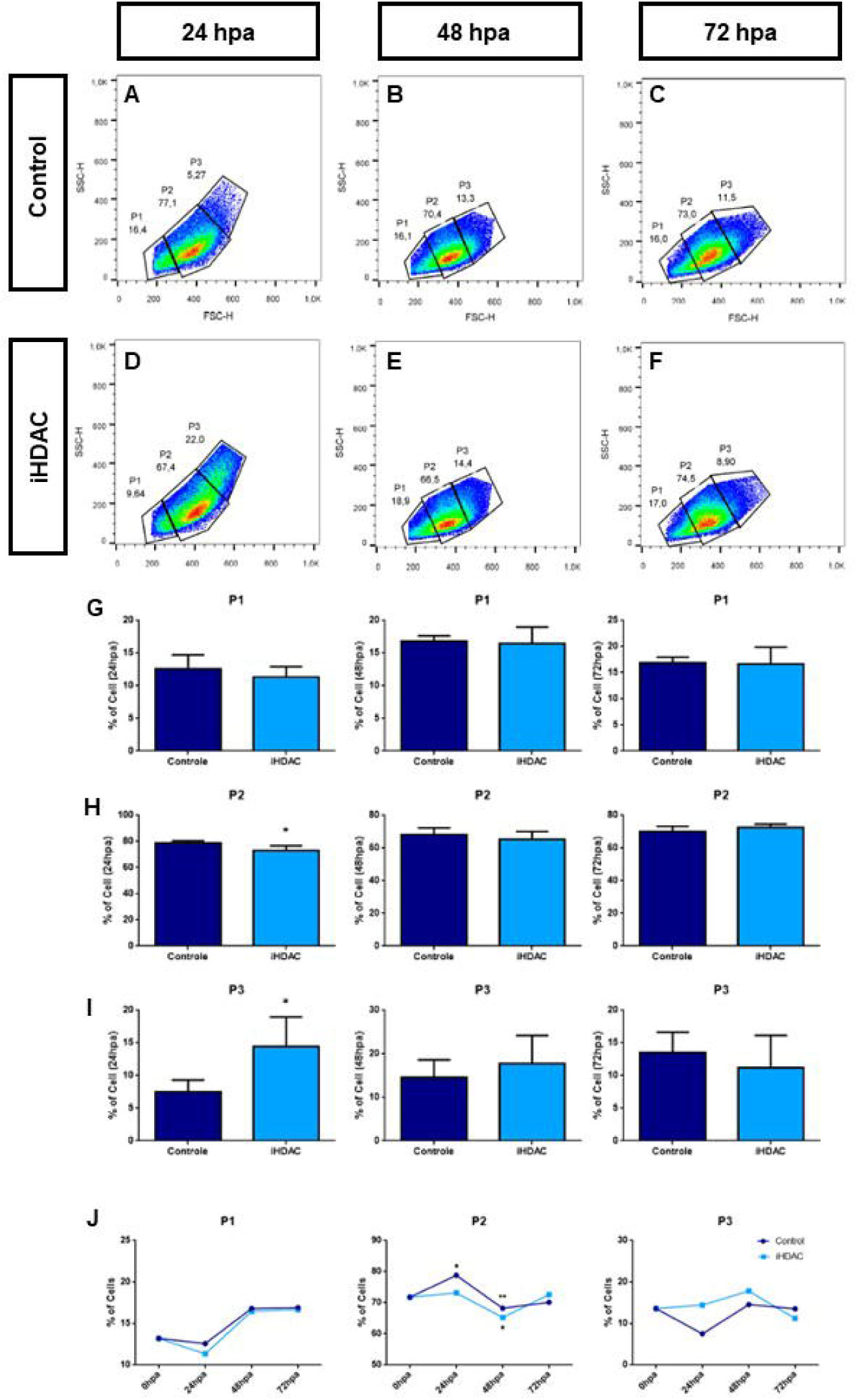
HDAC activity is necessary for the proper establishment of cellular dynamics in the regenerative bud at 24 hpa. Size (FSCH) and granularity (SSC-H) analysis of the cells present in the regenerative buds at 24 (A,D), 48 (B,E) and 72hpa (C,F). Three different cellular sub-sets were defined in accordance with the size and granularity where P1<P2<P3. Quantification of the percentage of cells in P1 (G), P2 (H) and P3 (I) along the time. Statistical analysis was performed using unpaired t-test, p <0.05. N = 3 (each biological replicate comprised 80 regenerative buds). (J) Analysis of the percentage of cells compared to “time zero” or steady state normal tails along the time. Statistical analysis was performed between a given time point and the point immediately before. Statistical analysis was performed using the Two-Way ANOVA test where the confidence interval is 95%. All analyzes were performed only using cells negative for the death label 7AAD (FL3-H). \**p*<0,05; \*\**p*<0,01.

Classical morphological analysis by May Grunwald-Giemsa showed the presence of acidophilic and basophilic structures in both control and iHDAC groups (supplementary Figure 1). To identify the presence of monocytes/macrophages among these cells we have used cytochemistry analysis for naphthyl esterase. In fact, we observed naphthyl esterase+ cells in both experimental groups in 24, 48 and 72 hpa (Supplementary Figure 2). In addition, we also examined the phagocytic activity of the cells present in the regenerative bud by neutral red uptake assay. In both experimental groups we were able to observe cells displaying phagocytic activity at 24, 48 and 72 hpa (Supplementary Figure 3).

We conclude that HDAC activity is necessary for the proper establishment of cellular dynamics in the regenerative bud at 24 hpa. In addition, we identified that cells displaying phagocytic activity with cytochemistry markers of monocytes/macrophages are components of the cellular population observed in the regenerative bud of *Xenopus* tadpoles.

### Myeloid cells are necessary for *Xenopus* tail regeneration in HDAC-activity dependent fashion

Next, we sought to demonstrate the myeloid origin of the cells above reported. Spib is one of the earliest genes to be expressed in the myeloid differentiation cascade at stage 17 while Xpox2, the ortholog of zebrafish mpox coding for myeloperoxidase and commonly expressed by neutrophils since embryogenesis in mammalians, is first detected in the anterior-ventral mesoderm at stage 19 and, at the tail-bud stage, can be found throughout the whole embryo (S. J. Smith et al., 2002). In fact, it has been shown, starting from stage 19, the existence of cellular sub-set whose phenotype is Spib^+^/Xpox2^+^, mostly representing granulocytes. In addition, mmp7 (matrilysin), a mettaloproteinase whose activity is linked to extra cellular matrix remodeling, has been described to be expressed at stage 18 in a punctate pattern in the aVBI (Harrison et al., 2004). MMP7^+^ cells represent phagocytic cells derived from Spib^+^ precursors such as macrophages (Costa et al., 2008) that lie in between the epidermis and mesoderm compatible with a pattern displayed by resident macrophages at stages 37/38 (Harrison et al., 2004). In fact, Spib morpholino injected embryos lack the expression of both downstream targets Xpox2 and mmp7 resulting in a lack of primitive myeloid cells (Costa et al., 2008).

We then hypothesized whether myeloid cells are necessary for *Xenopus* tail regeneration and whether HDAC activity is necessary to their behavior. To start addressing this question, we injected embryos with Spib anti-sense morpholinos at the 4 cells stage and, at the stage 40, the tadpoles were amputated, and the regeneration was scored at 72 hrs post amputation.

To access Spib knockdown we first performed an *in situ* hybridization for mmp7 mRNA at the neurula stage (stage 20). In fact, Spib morpholino injected embryos presented a substantial reduction in the mmp7 expression domain (Figure 3). Using two different concentrations of Spib morpholinos we consistently observed that Spib morpholino injected embryos present an impairment in tail regeneration when compared to scramble injected embryos. Spib knockdown led to 30% of non-regenerating tadpoles in the absence of considerable toxicity (Figure 3).

**Figure 3:**
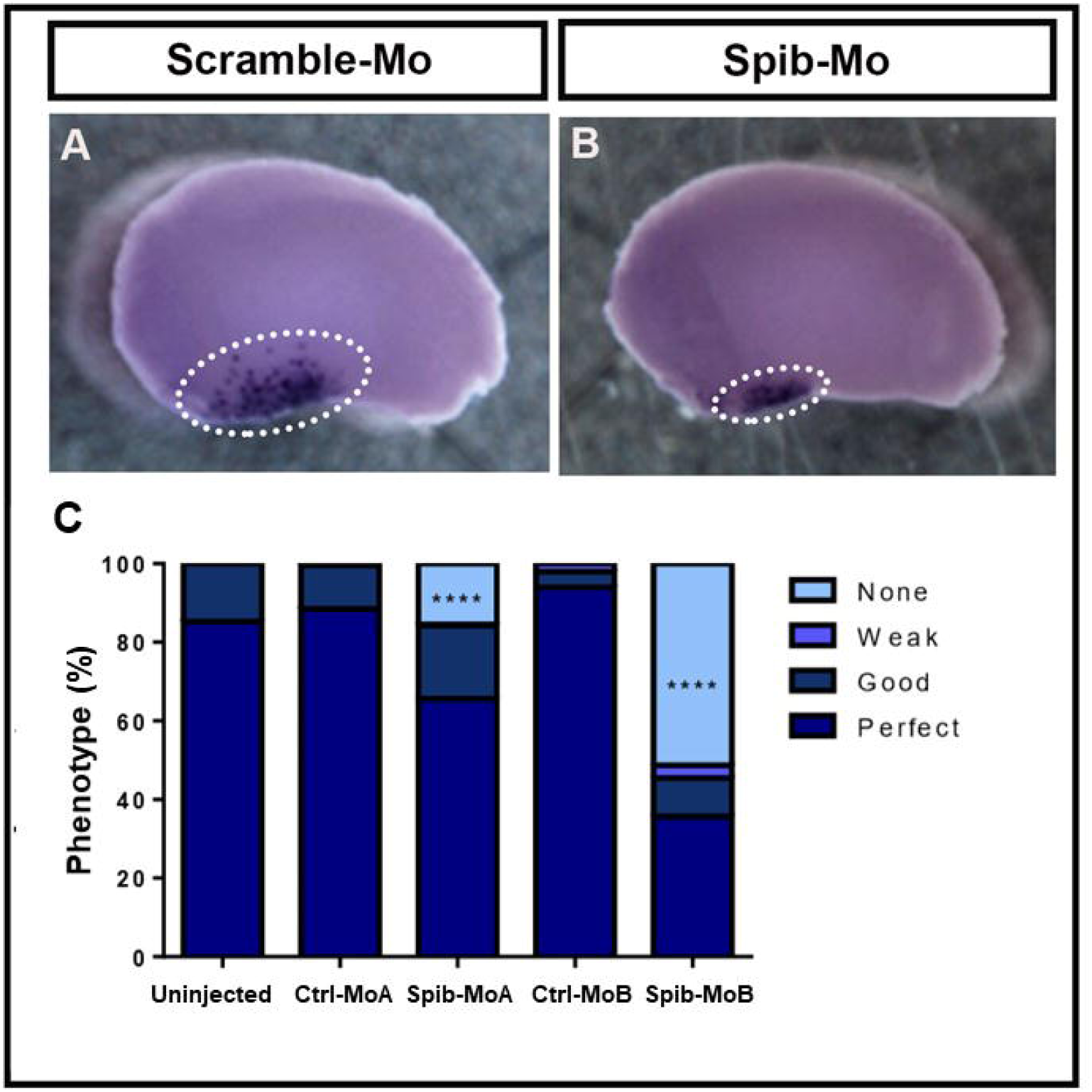
Spib is necessary for *Xenopus* tail regeneration in a dose dependent fashion. (A) Control-Morpholino (Ctrl-MoA) and (B) Spib-Morpholino (Spib-MoA) injected embryos were developed for *mmp7 in situ* hybridization. White dashed circle indicates the aVBI. The total number of viable amputated animals corresponds to: Ctrl-Mo A: 259; Spib-Mo A: 288; Ctrl-Mo B: 98 and Spib-Mo B: 153. Controls were compared to non-injected embryos, and each time of Spib-Mo injections was compared to their respective control. Statistical analyzes were performed using the chi-square test. **** *p*<0,0001.

Besides Spib, Xpox2 and mmp7, Smith et al described a myeloid-specific gene promoter that provided a molecular marker for myeloid cells at the early tail-bud stage. In this work Smith et al proposed that Ly-6/uPAR-related protein (XLURP-1) positive cells would be divided in two different sub-sets of myeloid cells: XLURP-1^+^/Xpox2^+^ and XLURP-1^-^/Xpox2^+^ suggesting the existence of at least two different cellular sub-sets of myeloid cells starting at the late tail-bud stage. On the other hand, despite displaying a similar punctate pattern of Xpox2, XLURP-1 transcripts were detected only starting from stage 24 onward. At stage 41 XLURP-1 positive cells present irregular shape suggesting that they may be resident phagocytic cells such as macrophages (S. J. Smith et al., 2002). In addition, XLURP-1 positive cells were found in the peripheral blood vessels suggesting a patrolling behavior.

Because Xpox2 and XLURP-1 also identifies myeloid populations in the developing embryo, we hypothesized whether XLURP-1 positive cells would be recruited into the regenerative bud, in a HDAC-activity dependent fashion, forming an infiltrate-like. LURP-GFP transgenic tadpoles were raised until stage 40 when were amputated and immediately placed in a solution containing iHDAC or DMSO for 24, 48 or 72hrs and at every time point, the number of GFP^+^ cells was counted/area of tissue. We observed at 24 hpa a consistent increase in the number of GFP^+^ /area in iHDAC treated tadpoles (Figure 4). In addition, we also quantified the endogenous levels of LURP mRNA by real time PCR at 24 and 72 hpa. Regardless we did not observe any change in transcripts levels at 24 hpa between control and iHDAC tadpoles, we detected an increase in LURP mRNA levels at 72 hpa in the iHDAC group (Supplementary Figure 4).

**Figure 4:**
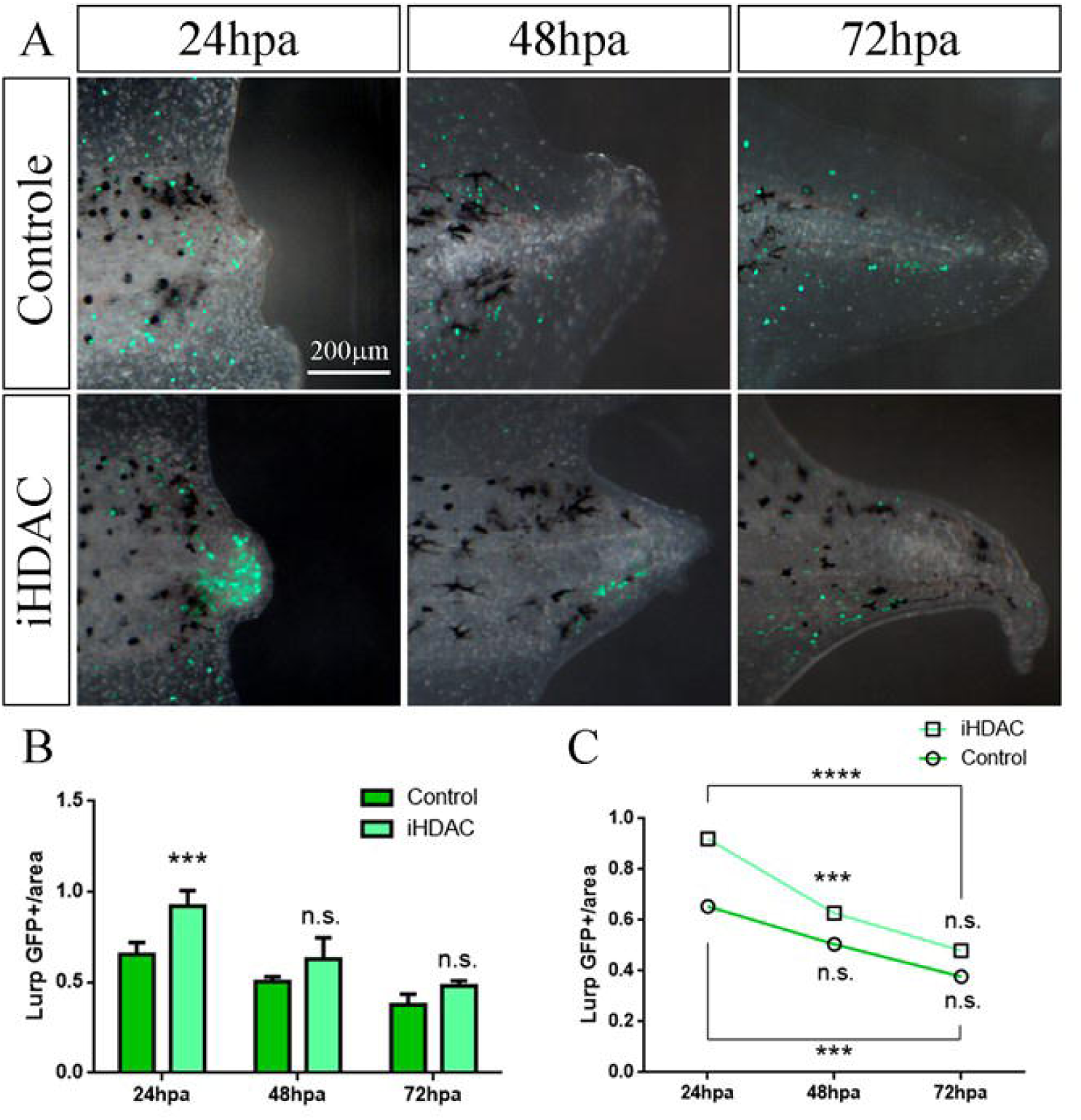
The absence of HDAC activity disrupts the pattern of LURP-GFP+ cell mobilization into the regenerative bud by 24hpa. (A) Fluorescence images show that the recruitment of LURP-GFP+ cells into the regenerative bud is disrupted upon iHDAC treatment when compared to control group (B) along the time (C). Quantification (B and C) was made taking into account the number of GFP+ puncta per area in µm2 of the regenerative bud (n ∼ 10 larvae/group; 5 repetitions). Two-Way ANOVA test. \*\*\**p*<0,001; \*\*\*\**p*<0,0001.

We conclude that LURP^+^ cells are necessary for tail regeneration and that their behavior is modulated by HDAC activity. We propose that LURP^+^ may be the same ones identified as the P3 sub-set, which displayed high size and relative granularity and may in turn, correspond to myeloid phagocytic cells.

### HDAC activity is necessary for the proper establishment of an inflammatory gene expression pattern during tail regeneration

Love et al have successfully accessed the initial inflammatory response following *Xenopus* tail amputation. In fact they described through *in situ* hybridization and sudan black staining, that mmp7 mRNA is up regulated as well as neutrophils, respectively, as early as 6 hrs post amputation, remaining for several days (Love et al., 2011). Here we asked whether HDAC activity would modulate the relative expression of mmp7, MPOX and spib during tail regeneration. To answer this question, we performed real time PCR to quantify the expression levels of this set of transcripts. In fact, at 24 hpa we observed a decrease in mmp7 mRNA expression levels, that was maintained at 72 hpa (Figure 5). On the other hand, we observed that spib and MPOX mRNA expression levels were increased when compared to control groups. These results argue in favor of an epigenetic control of the inflammatory response during tail regeneration. Given that MPOX is a myeloid marker for neutrophils, we interpret that HDAC activity blockade led to the persistence of the inflammatory response at 72 hpa, which is inversely proportional to the regenerative ability. In addition, increased levels of spib mRNA indicates that more undifferentiated myeloid cells are present in the regenerative bud if HDAC activity is suppressed. However, at this developmental stage, we are not able to separate myeloid cells form the first and the second wave of hematopoiesis. This hypothesis is in agreement with low levels of mmp7 mRNA expression levels which also argue in favor of a low phagocytic activity of myeloid cells and as a consequence, low inflammatory response.

**Figure 5:**
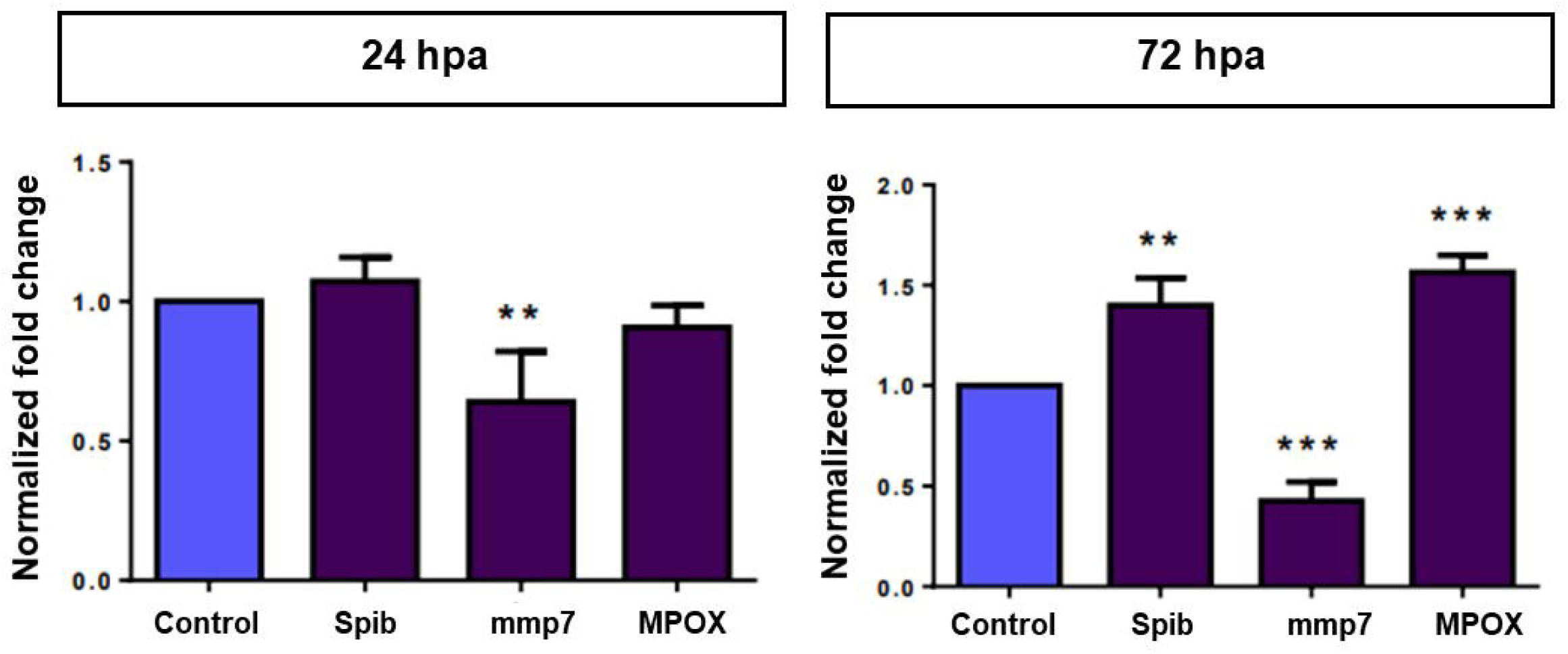
HDAC activity blockade disrupts the gene expression pattern of myeloid genes. Real time PCR for expression analysis of Spib, mmp7 and Mpox mRNAs (N=3; 50 tails/reaction). The endogenous housekeeping gene EF1α was used for normalization by the ΔΔCT method. Two-way ANOVA test with a 95% confidence interval was used. \*\*\**p*<0,001; \*\*\*\**p*<0,0001.

We conclude that HDAC activity is necessary for the establishment of transcriptional patterns compatible with the inflammatory response during tail regeneration.

### The inflammatory response is a key player during *Xenopus* tail regeneration and HDAC activity is necessary for its proper establishment

During tissue regeneration, acute inflammation has been shown as key player to trigger the main mechanisms that will drive regeneration. In fact, different works in zebra fish, salamander and mouse have addressed this question. To address whether the inflammatory response plays a functional/physiological role during *Xenopus* tail regeneration, we decided first to investigate the temporal dynamics of lipid droplets (LD) upon tail amputation. LD are osmiophilic subcellular components of leucocytes such as monocytes/macrophages and neutrophils that are involved during the inflammatory response. Upon the inflammatory stimulus, LD may increase either in size and in number and serves as a reservoir of lipid mediators and their enzymes, including 15-lipoxigenase (15-LOX)(Melo et al., 2011). As we have showed that HDAC activity blockade led to a disruption in the populational dynamics of leucocytes during tail regeneration, we decided to characterize the spatial and temporal dynamics of LD upon tail amputation and under the blockade of HDAC activity. For this, LD were visualized by conventional Oil Red staining and Osmiun tetroxide impregnation followed by light microscopy. We have shown that naphthyl esterase+ cells (monocytes/macrophages) are among the cellular components during tail regeneration. In addition, real time PCR also showed evidences for the presence of neutrophils (MPOX). These two cellular types are tight correlated to the inflammatory response and may represent the main source of LD in the context of tail regeneration. In fact, we were able to characterize the pattern of Oil Red+ vesicles distribution along the time in control and iHDAC tadpoles when compared to untreated, normal tails. It was possible to note that the number of Oil Red+ vesicles/area of tissue decreased along the time during normal tail development (Figure 6I, red line). When control and iHDAC groups were analyzed at 24 and 72 hpa, we observed that the overall dynamics was not disrupted, but the number of Oil Red+ vesicles/area of tissue was lower than the number observed for normal tails (Figure 6A,E,C,G,I). However we did not detected significant differences between control and iHDAC tadpoles at 24 and 72 hpa (Figure 6I). Using osmium impregnation we were able to detect multilocular cells at 24 hpa in semithin sections for electron microscopy preparations that were no longer observed at 72 hpa in control tadpoles (Figure 6B, B’, D). In iHDAC treated tadpoles we did not observe multilocular cells but instead, we observed osmiophilic vesicles ate 72 hpa that were not observed in control groups (Figure 6D,H). To better address whether LD, and their apparatus, would physiologically play a role during tail regeneration, we hypothesized that 15-LOX activity would be necessary for tail regeneration. To investigate this hypothesis we used baicalein to pharmacologically inhibit 15-LOX activity during tail regeneration. For this, amputated tails were placed in baicalein soon after amputation and aged until stage 47/48, when tadpoles were scored. In fact, 15-LOX blockade during tail regeneration led to tail regeneration impairment (Figure 6J). This result demonstrates that lipid mediators synthesized from LD play a role during *Xenopus* tail regeneration.

**Figure 6:**
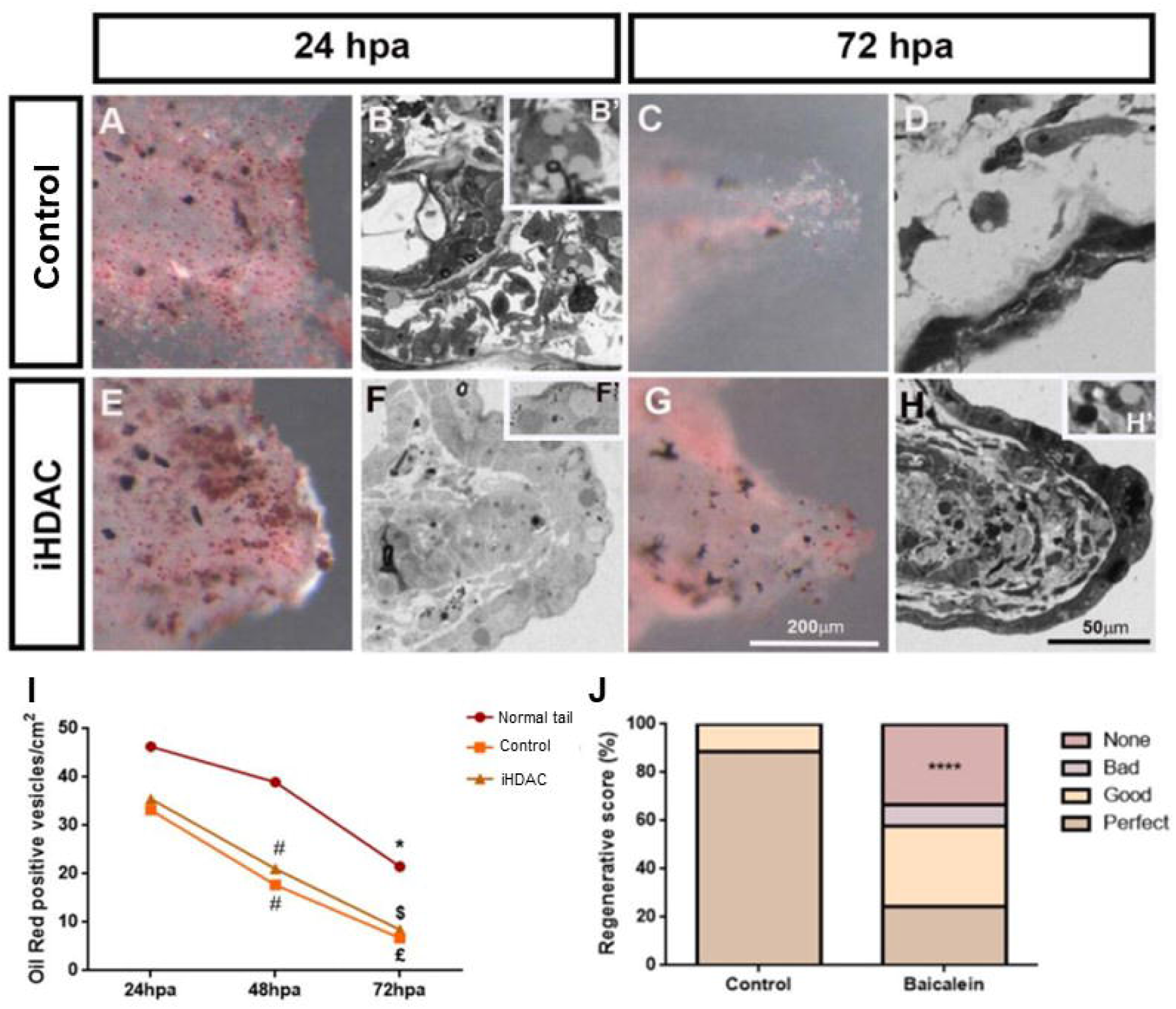
The inflammatory response is necessary for tail regeneration. Oil Red-O+ vesicles were observed in both control (A,C) and iHDAC treated tadpoles (E,G) at 24 and 72 hpa. (B) Comparisons among non-amputated, control and iHDAC treated tadpoles were made with the time point immediately before inside each one of these experimental groups. The quantification (B) was made taking into account the number of vesicles per area, in cm2, of the regenerative bud (N=10), using the Image J pluggin cell counter. The test used was the Two-Way ANOVA. *,#,$£ *p*<0,05. Semithin electron microscopy preparations show multilocular cells in control group (B,B’) that were not observed in iHDAC treated tadpoles (F,F’) at 24 hpa. At 72 hpa iHDAC treated tadpoles presented osmiophilic vesicles (H,H’).

We conclude that the inflammatory response is a key player during *Xenopus* tail regeneration and that HDAC activity is necessary for its proper establishment.

## Discussion

In the present paper we aimed to characterize the epigenetic control HDAC-dependent of tissue and organ regenerative capacity using *Xenopus* tail as experimental model. Regardless it is already well established in the literature that the acute inflammatory response, through an injury, exerts a protective effect on the organism (Kyritsis et al., 2012; Mescher, Neff, & King, 2017), tractable strategies to modulate inflammation favoring tissue and organ regeneration are still lacking.

Similarly to mammals, the hematopoiesis in amphibians generates myeloid cells with critical physiological roles both during embryonic and adult life. In both animals the first wave of differentiation gives rise to myeloid cells that will spread in the embryo and play crucial roles during morphogenesis and regeneration (Agricola et al., 2016; Costa et al., 2008). We showed that in the absence of HDAC activity the number of LURP-GFP^+^ cells in the regenerative bud was increased without a consistent increase in the levels of endogenous mRNA levels of LURP. Interpreting this result, we found a stinking similar pattern observed in mice upon iHDAC treatment of myeloid precursors at 24 hpa (Cabanel et al., 2015; Cabanel et al., 2019). While control group displayed a lower number of LURP-GFP+ cells, these cells might express high levels of this protein being classified as LURP-GFP^high^ sub-set (pro-inflammatory monocytes). In contrast, because iHDAC treated tadpoles presented increased number of LURP-GFP+ cells, they may belong to a different sub-set of cells, which in turn will display low levels of LURP-GFP+ being classified as LURP-GFP^low^ (patrolling/anti-inflammatory/pro-resolutive monocytes). This may explain why despite a higher number of total LURP-GFP+ cells, iHDAC treated tadpoles did not present a consistent increase in LURP mRNA levels at this time point. This scenario is also compatible with low levels of mmp7 mRNA expression, which indicates low inflammatory activity. Taken together these results indicates that iHDAC treatment impaired the inflammatory response at 24 hpa compromising, in this way, the outcome of the main regenerative events that will follow.

In fact, this interpretation is in agreement with the results obtained at 72 hpa. At this time point regardless the number of LURP-GFP+ cells is the same in control and iHDAC treated tadpoles, LURP mRNA expression levels are increased in iHDAC tadpoles. This result indicates that in iHDAC treated tadpoles, LURP-GFP+ cells belong to the LURP-GFP^high^ sub-set, compatible with the persistence of the inflammatory response at 72 hpa correlated to a lack of regeneration. This rational is supported by the fact that at this time point MPOX mRNA expression levels are increased along with increased levels of Spib mRNA (more undifferentiated cells) and decreased levels of mmp7 mRNA expression (less active phagocytic cells)(Harrison et al., 2004). Taken together these results argue in favor of a disruption in the assemble of the inflammatory response in the absence of HDAC activity. This interpretation is also aligned with the fact that mRNA levels of IL11, a interleukin involved in tissue repair and regeneration, are increased at 72 hpa (Tsujioka, Kunieda, Katou, Shirahige, & Kubo, 2015), indicating that at this time point the inflammation must to be resolved, which is not the case if HDAC activity is inhibited.

It has been documented that LURP and MPOX colocalize in myeloid cells and that give rise to mmp7+ macrophages (S. J. Smith et al., 2002). Our results agree with this report given that both transcripts present the same pattern of expression along with Spib at 24 and 72 hpa. In fact, in iHDAC treated tadpoles these transcripts were found upregulated, if compared to control groups, supporting the view that myeloid cells are the main target for the epigenetic control through HDAC activity blockade during tail regeneration. In addition, Love et al have shown that myeloid cells are observed in the regenerative bud upon tail amputation at 6 hpa along with a clear dynamic of mmp7 expression, that is observed also at 6 hpa, is sustained at 24 hpa but diminished by 48 and 72 hpa (Love et al., 2011). In contrast, when HDAC activity was inhibited, the mRNA levels of mmp7 remained stable in 24 and 72 hpa, showing that HDAC activity is required to the physiological role of myeloid phagocytic cells, mostly resident macrophages. This result is also in agreement with the findings reported by Harrison and colleagues showing that mmp7 co-localizes with MPOX (Harrison et al., 2004).

Another important piece of evidence that corroborates the persistence of the inflammatory response at 72 hpa in iHDAC treated tadpoles, comes from the analysis of LD dynamics. These subcellular organelles in immune cells are key players in the assemble of the inflammatory response because they can store arachidonic acid to the synthesis of eicosanoids (Bozza, Magalhães, & Weller, 2009; Weller, Ryeom, Picard, Ackerman, & Dvorak, 1991) upon the inflammatory stimulus by enzymes such as 15-LOX, leading to the synthesis of pro-resolving mediators as Resolvins (Serhan, 2014). Although we did observe any overall difference in the profile of LD among normal tail, control and iHDAC treated tadpoles during tail regeneration using Oil Red, we observed the presence of osmiophilic LD in iHDAC treated tadpoles at 72 hpa. The presence of these organelles indicates that the inflammatory response is still active at 72 hpa in iHDAC treated tadpoles if compared to control group. The persistence of inflammatory mediators would contribute to regeneration impairment.

Thus, our working model suggest two different time points in which HDAC activity is necessary for tail regeneration. In a first moment, at 24 hpa, HDAC blockade dampens the inflammatory response from the onset, favoring patrolling/anti-inflammatory monocytes (LURP^low^) in detriment of pro-inflammatory monocytes (LURP^high^) (Figure 7). This is consistent with low expression levels of mmp7 at this time point. In a second moment, as an attempt of the organism to proper assemble the inflammatory response, more myeloid progenitor Spib, LURP and MPOX positive cells are recruited into the regenerative bud, as an infiltrate-like. However, HDAC activity blockade leads to the persistence of inflammatory myeloid cells MPOX and LURP^high^ cells at 72 hpa resulting in loss of the regenerative ability (Figure 7).

**Figure 7:**
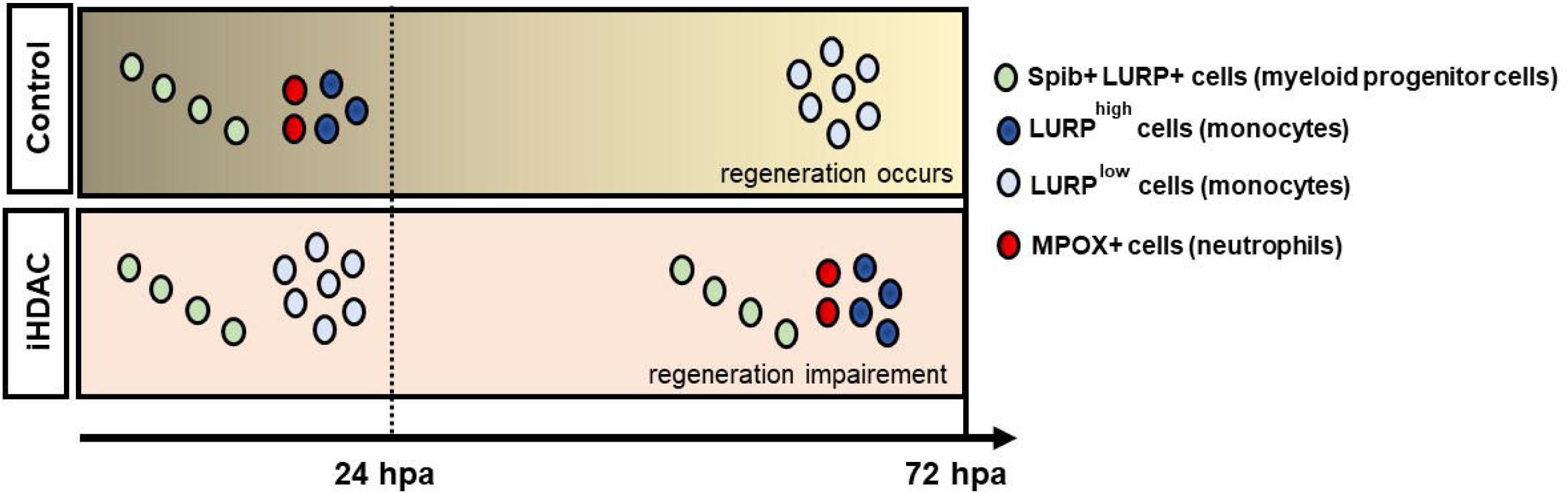
Working model. At 24 hpa, while HDAC blockade dampens the inflammatory response from the onset, favoring the differentiation of Spib+ cells (green dots) in patrolling/anti-inflammatory monocytes (LURP^low^; light blue dots) in detriment of pro-inflammatory monocytes (LURP^high^; dark blue dots), control animals present pro-inflammatory monocytes (LURP^high^; dark blue dots) as well as neutrophils (red dots). At 72 hpa, as an attempt of the organism to proper assemble the inflammatory response, more myeloid progenitor cells Spib, LURP and MPOX positive cells are recruited into the regenerative bud of iHDAC treated tadpoles, as an infiltrate-like. However, HDAC activity blockade leads to the persistence of inflammatory myeloid cells MPOX and LURP^high^ cells at 72 hpa resulting in loss of the regenerative ability.

Our results show for the first time in the literature that Epigenetic mechanisms HDAC-dependent modulate the behavior of myeloid cells during tissue and organ regeneration. Because iHDACs have been widely studied in the literature and some iHDACs have also been approved for use in humans, we propose that the use of this class of inhibitors may be converted in a tractable strategy in translational studies to better understand their potential in clinical contexts.

## Material and Methods

### Animals

The larvae of *Xenopus laevis* were obtained in collaboration with Prof. José Garcia Abreu (ICB-UFRJ). After fertilization, the embryos were aged to stage 40 (approximately 3 days after fertilization -at 23°C or 7 days at 14°C) for regeneration assays. Larvae were maintained in Barth culture medium or Steinberg. Stage 40 *Xenopus* larvae were anesthetized with 2mM Tricaine in Barth or Steinberg 1x medium, and amputated at the final third of the tail, as described in (A.-S. Tseng et al., 2011), and separated into two experimental groups: Control (larvae maintained in DMSO culture medium) or iHDAC group (culture medium containing 25nM of the HDAC pharmacological activity inhibitor Tricostatin A - TSA). Adult animals were kept in accordance with CEUA 152/13 authorization. The larvae submitted to the experiments were kept according to CEUA 045/19 authorization.

### Molecular Biology

#### Primer synthesis

The primers used for the RT-PCR reaction were designed using the Primer Blast program from the National Center for Biotechnology Information (NCBI) web page (https://www.ncbi.nlm.nih.gov/tools/primer-blast/) and purchased form Integrated DNA Technologies (IDT).

EF1α: F: GCCCCAATATGCCTTGGTT R: ATGCAGTCAAGAGCTTCCAG

Spib: F: AAGGAATTGCTGGCTCACCG R: ACTGTCAAACTGATACGTCAGC

LURP1: F: GCTGAAGCTTTGAAGTGTCG R: AAGCACAGAGTGGAATCTCG

Mpo (Pox2): F: TCCTTCACAGGGGAGTGTAA R: TCTCTGTCCATCCTTTGGGA

Mmp7: F: CCAGCTGACATACAGCATTG R: TGCGTCTCCAGAGGAAATTG

### mRNA extraction

Larvae at stage 40/41 were anesthetized, amputated and divided between the Control or iHDAC groups. At 24hpa and 72hpa, 50 larvae of each condition were anesthetized and had their regenerative buds dissected in a petri dish. The buds were placed in Eppendorf tubes in ice, containing the culture medium and the anesthetic. At the end of the dissection, the tubes were rapidly centrifuged to concentrate the buds allowing the medium to be removed. 50µL of Trizol (Ambion) was added in the tubes to be homogenized. An additional 450µL of Trizol was added and RNA extraction proceeded as manufacturer’s specifications. If extraction could not be done at the same day, the tubes were stored in a freezer at 80°C with Trizol. After extraction, RNA was suspended in 21µL of RNase free water and 1µL of each sample was analyzed using NanoDrop followed by storage at -80°C until use.

### cDNA synthesis

cDNA synthesis was performed using the enzyme Multiscribe (Applied Biosystems). The total amount of mRNA required for cDNA synthesis is 1μg according to manufacturer’s instructions. At the end of the reaction, 1µL of each sample had its concentration measured in Nano Drop followed by storage at -20°C until use.

### Real Time PCR Chain Reaction

The reaction was performed using SYBR™ Green PCR Master Mix (Applied Biosystems), 100ng cDNA and 500nM of each primer. We use the following conditions for the 40-cycle reaction on the Quant Studio TM 7 Flex System (Thermo Fisher): a. 10 minutes at 95 ° C b. 15 seconds at 95 ° C c. 1 minute at 60 ° C d. 15 seconds at 95 ° C e. 1 minute at 60 ° C f. 15 seconds at 95°C. Relative quantification was calculated using the ΔΔCt method using endogenous EF1-α. Statistical analyzes were performed using the GraphPad Prism 6 software by the two-way ANOVA test.

### Flow Citometry

Larvae at stage 40 were anesthetized, amputated and divided between Control or iHDAC groups (except for experimental time 0hpa). At 0, 24, 48 and 72hpa, 80 larvae of each condition were anesthetized and their regenerative buds were dissected in a petri dish. The buds were placed in Eppendorf tubes on ice, containing the culture medium and anesthetic. At the end of the dissection, the tubes were rapidly centrifuged to concentrate the buds at the bottom of the tubes, allowing the medium to be removed. 1mL collagenase (SigmaAldriech) was added for tissue dissociation, followed by incubation for 30min at 37 ° C. The samples were filtered using 40µm filters (Cell Strainer - Falcon®) and transferred to Falcon® tubes. 6mL of 1x PBS was added and the samples were centrifuged for 10min at 4 ° C, 259rcf. The pellet was resuspended in 2mL of 1x PBS and centrifuged to clean the sample. The pellet was resuspended in 1mL of 1x PBS and a 10µl aliquot was taken for Neubauer chamber analysis. Another 100 µL aliquot was taken and used for centrifugation in Cytospin 4 (Thermo scientific) for 3min, room temperature, 14rcf. The remaining samples were centrifuged and resuspended in 300µL of 1x PBS on ice for analysis on the FACS Calibur flow cytometer using CellQuest software, where a total of 100,000 events were collected from each sample. The viability of each sample was seen by incubating 7AAD (BD-Pharmingen) for 10min on ice. Only viable cells were analyzed using the FlowJo program, and the results were expressed as a percentage of cells distributed in the delimited regions.

### May-Grünwald Giemsa Staining

Cytocentrifugation slides were immediately fixed in methanol at room temperature overnight. After fixation, the slides were subjected to the filtered May-Grünwald (Merck) dye solution diluted 1: 2 in distilled water for 10 min at room temperature. The slides were then washed in distilled water and subjected to the filtered Giemsa dye solution (Merck) diluted 1:20 in distilled water for 20 min at room temperature. The slides were washed in distilled water and after drying were sealed with Entellan (Merck). The images were taken through the Axioplan (Carl Zeiss) brightfield microscope using AxioVision (Carl Zeiss) software.

### α-naphthyl acetate (NSE) reaction

According to the methodology, α-naphthyl acetate is enzymatically hydrolyzed, releasing a compound that attaches to a diazonium compound, forming black deposits at nonspecific esterase activity sites. This enzyme is mainly detected in monocytes, macrophages and histocytes and is usually absent in granulocytes. Slides from cytocentrifugations were immediately fixed in CAF buffer (Citrate - supplied by Sigma Kit 91A; Acetone - Proquimics; Formaldehyde - Proquimics) and processed according to Kit 91A (Sigma) protocol for monocyte detection. The images were taken using the Axioplan (Carl Zeiss) brightfield microscope using AxioVision (Carl Zeiss) software.

### Neutral red

Stage 40 larvae were anesthetized, amputated and divided between the Control or iHDAC groups, both containing 5µg / ml of the non-toxic neutral red dye. Incubation with this non-toxic dye lasted 6h, followed by a culture medium wash and incubation with medium containing only DMSO or TSA. The culture media were changed once a day, at 24, 48 and 72hpa collection points, when the larvae were anesthetized, euthanized and fixed in 1X MEMFA for 2 hours at room temperature (RT) or overnight at 4 ° C. These larvae were kept in 1x PBS and photographed in fluorescence stereoscope (Leica) using the red filter (DSRed) to identify neutral red particles.

### Recruitment of xLURP-GFP cells

xLURP-GFP transgenic larvae at stage 40 were anesthetized, amputated and divided between the Control or iHDAC groups (n = 50). At 24, 48 and 72hpa 10 larvae were randomly selected and anesthetized to be photographed under a Nikon fluorescence microscope. This experiment was repeated five times. The images were analyzed by ImageJ software, where the regenerated area was delimited and measured. Using the Cell Counter pluggin, we quantify GFP + cells only within the delimited area. Thus, the ratio of GFP + cells to the regenerated area was analyzed. The average of these ratios was plotted in the GraphPad Prism 6 program. Two-Way ANOVA test was used.

### Spib knockdown

Mismatch and Spib morpholinos were purchased from Gene Tools based on (Agricola et al., 2016; Costa, Soto, Chen, Zorn, & Amaya, 2008b; Smith & Mohun, 2011). According to the manufacturer’s recommendations, 300µL of RNase-free water was added resulting in a 1mM solution. This stock solution was stored in aliquots in at -20 ° C until use. The sequences used are listed below and the control morpholino red bases (Ctrl-Mo) correspond to changes from the morpholino Spib (Spib-Mo) bases.

Spib-Mo (Spib-a-e1i1-Mo): 5’AAA TAC AGT CCA TCA CTT ACC GGA G

Ctrl-Mo (Mismatch-ctrl): 5’AAA TAG ACT GCA TCA CTT AGC CGA G

Morpholino injection was performed on both dorsal blastomers of stage 3 embryos (4 cells). We have used two injection times, 70 (Spib-morpholino A) and 100 milliseconds (Spib-morpholino B), respectively. After injection the embryos were kept in 0.1x MMR culture medium and aged to stage 22/23 (n = 15), when they were fixed for in situ hybridization, or until stage 40, when they underwent amputation of the final third of the tail having its regeneration evaluated after 72hpa.

### *in situ* hybridization

Fifteen embryos injected with 4µg / µL Spib-MO at stage 22/23 were randomly selected and fixed in 1X MEMFA for 2h at room temperature. Embryos were progressively dehydrated in Methanol / PBST (Tween 0.1% in 1X PBS) (25%, 50%, 75% and 100%) for 5 min each and stored at 4 ° C. For *in situ*, embryos were progressively rehydrated in PBST / Methanol (25%, 50%, 75% and 100%) and prehybridized for 1h at 60 ° C followed by incubation with the mmp7 probe overnight at 60°C. Next, 3 washes of 20 min each were made in sodium citrate buffer (2x SSC), followed by 2 washes in 0.2 x SSC for 30 min each, at 60 ° C. Subsequently, incubation with maleic acid / NaCl (MAB) was performed for 10 min at room temperature, followed by blocking reaction (MAB + 2% Block reagent) for 1h at room temperature. The anti-digoxigenin antibody (α-DIG) incubation was performed at a 1: 3000 dilution in blocking solution for 4h at room temperature, followed by 3 washes with MAB and incubation overnight at 4 ° C. Next, 3 washes in MAB for 20 min at room temperature were done, followed by 2 washes in alkaline phosphate buffer (AP Buffer) and incubation with BM-Purple for 4h at 37 ° C for chemical developing. A 2x SSC wash and incubation with a bleaching solution containing 2.5% SSC20X, 5% formamide, 4% hydrogen peroxide 30% was performed. The embryos were washed twice with 1X PBS and transferred to PBST for further image processing. Embryos were dehydrated in ethanol and stored at -20 ° C.

### Oil Red-O staining

Larvae at stage 40 were anesthetized, amputated and divided between the Control or iHDAC groups. At 24hpa and 72hpa, 10 larvae of each condition were anesthetized and fixed in MEMFA (0.1M MOPS pH7.4; 2nM EGTA; 1nM MgSO4; 3.7% Formaldehyde) for 2h at room temperature. The larvae were stored in PBS at 4°C for no more than one week. Larvae were placed in 12-well plates and incubated with 0.2% fresh Oil Red solution in Isopropanol (Merck) for 15 min. After three successive washes with 1X PBS the larvae were photographed in Leica M205 FA stereoscope. This procedure was repeated 3 times. To quantify the number of oil red+ vesicles, the Image J program was used through the “Cell Counter” plugging. The ratio of the sum of oil red+ vesicles to the sum of the areas was used for statistical analysis using the Two-Way ANOVA test by GraphPad Prism 6.

### Semi-thin sections for electron microscopy

Stage 40 larvae were anesthetized, amputated and divided between the Control or iHDAC groups. At 24 and 72hpa, 5 larvae of each condition were anesthetized and their regenerative buds were dissected in a petri dish. The buds placed in Eppendorf tubes on ice, containing the culture medium and anesthetic. At the end of the dissection, the tubes were rapidly centrifuged to concentrate the buds allowing the medium to be removed. 1mL of fixative solution (4% PFA + 2% glutaraldehyde (pH 7.6) in 0.1M phosphate buffer) was added and the tubes were kept at 4 ° C until use.

Day 1: after fixation, samples were washed in 0.1M cacodylate buffer pH 7.4 (three 5-minute washes) and fixed in 1% osmium tethoxide + potassium ferrocyanide 0, 8% and 5 nM calcium chloride in 0.1 M cacodylate buffer pH 7.4 for approximately 60 to 90 minutes, being kept in the dar. After this step, the samples were washed again in 0.1 M cacodylate pH 7.4 buffer (three 5 minute washes each) and in distilled water (1 5 minute wash) and finally placed in 1% aqueous uranyl acetate solution. in the absence of light, all night under shaking. Day 2: Samples were washed three times in 0.1 M cacodylate pH 7.4 buffer, and dehydrated in a gradual 30%, 50%, 70%, 80%, 90% acetone (two washes of seven minutes each at each concentration) and 100% (two washes of fifteen minutes each). Then the material was infiltrated into a 1: 1 mixture of 100% acetone and resin (polybed 812) while stirring overnight. Day 3: Samples were infiltrated into pure resin and kept overnight for stirring. Day 4: Finally the materials were included in silicone molds using 100% resin and baked at 60 ° C for 48 hours for resin polymerization. After polymerization, each resin block was promptly debased and made into a trapezoidal pyramid to remove excess resin, and 500 nm semifinely cross-sections were made on an ultramicrotome (MT-600-XL-RMC, inc), placed in glass slides and stained with 1% aqueous toluidine blue solution. Semi-thin sections were photographed under Axioplan (Carl Zeiss) brightfield microscope using AxioVision (Carl Zeiss) software.

### Baicalein assay

Stage 40 larvae were anesthetized, amputated and divided between the Control or Baicalein groups. As it is a flavonoid with no references for use in *Xenopus laevis* larvae, it was necessary to perform toxicity tests. We used the concentrations of 10µM, 5µM, 4µM, 2.5µM and 1µM, diluted in Barth 1X culture medium. The concentration of 2.5µM presented phenotypes without affecting larval viability. The animals were kept at 23°C and protected from light as recommended by the manufacturer, and the medium was replaced once a day. The larvae were aged up to 72hpa, when they were anesthetized for regeneration score analysis.

## Acknowledgements

We would like to acknowledge Grasiella Ventura Matioszek for helpful support in Confocal Microscopy and Funding Agencies FAPERJ, CNPq and CAPES, for Nathalia Pentagna’s PhD Fellowship.

## Competing Interests

The authors declare no competing interests.

## Figure legends

**Supplementary Figure 1: Morphological characterization of cellular components of regenerative buds along the time.** May-Grünwald/Giemsa staining reveals eosinophilic and acidophilic structures present in the cellular components of regenerative buds in both control (A,C,E) and iHDAC treated tadpoles (B,D,F).

**Supplementary Figure 2:** Identification of myeloid cells by Naphthyl esterase reaction. At all experimental times it was possible to identify cells with black deposits, characteristic of the reaction of esterases present in myeloid cells, in both control (A,C,E) and iHDAC treated tadpoles (B,D,F). (A’) Positive reaction control performed in myeloid cells from bone marrow preparation from C57/BL6 mice. (E’,F’) Macrophage cells.

**Supplementary Figure 3: Phagocytic cells are cellular components of regenerative buds.** Upon larvae incubation with non-toxic neutral red for 6h after amputation, phagocytic cells were detected in both control (A,C,E) and iHDAC treated tadpoles (B,D,F) along the time. Brightfield images (A-F). Neutral red fluorescence (A’-F ’). Overlapping images (A’ ’-F’ ’). Anterior is to the left; posterior is to the right; dorsal is up and ventral is down.

**Supplementary Figure 4: HDAC activity is necessary for LURP mRNA expression pattern.** Real-time PCR for LURP mRNA levels at 24 and 72 hpa. The endogenous housekeeping gene EF1α was used for normalization by the ΔΔCT method. (N=3; 50 buds/ sample). Two-way ANOVA. \*\**p*<0,001.

## Notes

### Competing Interest Statement

The authors have declared no competing interest.

## References

Adams, D. S., Masi, A., & Levin, M. (2007). H+ pump-dependent changes in membrane voltage are an early mechanism necessary and sufficient to induce Xenopus tail regeneration. Development, 134(7), 1323–1335. DOI: 10.1242/dev.02812

Agricola, Z. N., Jagpal, A. K., Allbee, A. W., Prewitt, A. R., Shifley, E. T., Rankin, S. A., Kenny, A. P. (2016). Identification of genes expressed in the migrating primitive myeloid lineage of Xenopus laevis. Dev Dyn, 245(1), 47–55. DOI: 10.1002/dvdy.24314

Aurora, A. B., Porrello, E. R., Tan, W., Mahmoud, A. I., Hill, J. A., Bassel-Duby, R., Olson, E. N. (2014). Macrophages are required for neonatal heart regeneration. J Clin Invest, 124(3), 1382–1392. DOI: 10.1172/JCI72181

Beane, W. S., Morokuma, J., Lemire, J. M., & Levin, M. (2013). Bioelectric signaling regulates head and organ size during planarian regeneration. Development, 140(2), 313–322. DOI: 10.1242/dev.086900

Beck, C. W., Christen, B., & Slack, J. M. (2003). Molecular pathways needed for regeneration of spinal cord and muscle in a vertebrate. Dev Cell, 5(3), 429–439.

Bozza, P. T., Magalhães, K. G., & Weller, P. F. (2009). Leukocyte lipid bodies Biogenesis and functions in inflammation. Biochim Biophys Acta, 1791(6), 540–551. DOI: 10.1016/j.bbalip.2009.01.005

Cabanel, M., Brand, C., Oliveira-Nunes, M. C., Cabral-Piccin, M. P., Lopes, M. F., Brito, J. M., Carneiro, K. (2015). Epigenetic Control of Macrophage Shape Transition towards an Atypical Elongated Phenotype by Histone Deacetylase Activity. PLoS One, 10(7), e0132984. DOI: 10.1371/journal.pone.0132984

Cabanel, M., da Costa, T. P., El-Cheikh, M. C., & Carneiro, K. (2019). The epigenome as a putative target for skin repair: the HDAC inhibitor Trichostatin A modulates myeloid progenitor plasticity and behavior and improves wound healing. J Transl Med, 17(1), 247. DOI: 10.1186/s12967-019-1998-9

Chen, X. D., & Turpen, J. B. (1995). Intraembryonic origin of hepatic hematopoiesis in Xenopus laevis. J Immunol, 154(6), 2557–2567.

Ciau-Uitz, A., Liu, F., & Patient, R. (2010). Genetic control of hematopoietic development in Xenopus and zebrafish. Int J Dev Biol, 54(6-7), 1139–1149. DOI: 10.1387/ijdb.093055ac

Ciau-Uitz, A., Walmsley, M., & Patient, R. (2000). Distinct origins of adult and embryonic blood in Xenopus. Cell, 102(6), 787–796. DOI: 10.1016/s0092-8674(00)00067-2

Costa, R. M., Soto, X., Chen, Y., Zorn, A. M., & Amaya, E. (2008). spib is required for primitive myeloid development in Xenopus. Blood, 112(6), 2287–2296. DOI: 10.1182/blood-2008-04-150268

Godwin, J. W., Pinto, A. R., & Rosenthal, N. A. (2013). Macrophages are required for adult salamander limb regeneration. Proc Natl Acad Sci U S A, 110(23), 9415–9420. DOI: 10.1073/pnas.1300290110

Harrison, M., Abu-Elmagd, M., Grocott, T., Yates, C., Gavrilovic, J., & Wheeler, G. N. (2004). Matrix metalloproteinase genes in Xenopus development. Dev Dyn, 231(1), 214–220. DOI: 10.1002/dvdy.20113

Ho, D. M., & Whitman, M. (2008). TGF-beta signaling is required for multiple processes during Xenopus tail regeneration. Dev Biol, 315(1), 203–216. DOI: 10.1016/j.ydbio.2007.12.031

Kau, C. L., & Turpen, J. B. (1983). Dual contribution of embryonic ventral blood island and dorsal lateral plate mesoderm during ontogeny of hemopoietic cells in Xenopus laevis. J Immunol, 131(5), 2262–2266.

Kyritsis, N., Kizil, C., Zocher, S., Kroehne, V., Kaslin, J., Freudenreich, D., Brand, M. (2012). Acute inflammation initiates the regenerative response in the adult zebrafish brain. Science, 338(6112), 1353–1356. DOI: 10.1126/science.1228773

Lane, M. C., & Sheets, M. D. (2002). Primitive and definitive blood share a common origin in Xenopus: a comparison of lineage techniques used to construct fate maps. Dev Biol, 248(1), 52–67. DOI: 10.1006/dbio.2002.0717

Lane, M. C., & Smith, W. C. (1999). The origins of primitive blood in Xenopus: implications for axial patterning. Development, 126(3), 423–434.

Levin, M. (2007). Large-scale biophysics: ion flows and regeneration. Trends Cell Biol, 17(6), 261–270. DOI: 10.1016/j.tcb.2007.04.007

Levin, M. (2014). Endogenous bioelectrical networks store non-genetic patterning information during development and regeneration. J Physiol, 592(11), 2295–2305. DOI: 10.1113/jphysiol.2014.271940

Love, N. R., Chen, Y., Bonev, B., Gilchrist, M. J., Fairclough, L., Lea, R., Amaya, E. (2011). Genome-wide analysis of gene expression during Xenopus tropicalis tadpole tail regeneration. BMC Dev Biol, 11, 70. DOI: 10.1186/1471-213X-11-70

Medvinsky, A., & Dzierzak, E. (1996). Definitive hematopoiesis is autonomously initiated by the AGM region. Cell, 86(6), 897–906. DOI: 10.1016/s0092-8674(00)80165-8

Melo, R. C., D’Avila, H., Wan, H. C., Bozza, P. T., Dvorak, A. M., & Weller, P. F. (2011). Lipid bodies in inflammatory cells: structure, function, and current imaging techniques. J Histochem Cytochem, 59(5), 540–556. DOI: 10.1369/0022155411404073

Mescher, A. L., Neff, A. W., & King, M. W. (2017). Inflammation and immunity in organ regeneration. Dev Comp Immunol, 66, 98–110. DOI: 10.1016/j.dci.2016.02.015

Morales, R. A., & Allende, M. L. (2019). Peripheral Macrophages Promote Tissue Regeneration in Zebrafish by Fine-Tuning the Inflammatory Response. Front Immunol, 10, 253. DOI: 10.3389/fimmu.2019.00253

Paredes, R., Ishibashi, S., Borrill, R., Robert, J., & Amaya, E. (2015). Xenopus: An in vivo model for imaging the inflammatory response following injury and bacterial infection. Dev Biol, 408(2), 213–228. DOI: 10.1016/j.ydbio.2015.03.008

Petrie, T. A., Strand, N. S., Yang, C. T., Tsung-Yang, C., Rabinowitz, J. S., & Moon, R. T. (2014). Macrophages modulate adult zebrafish tail fin regeneration. Development, 141(13), 2581–2591. DOI: 10.1242/dev.098459

Rollins-Smith, L. A., & Blair, P. (1990). Contribution of ventral blood island mesoderm to hematopoiesis in postmetamorphic and metamorphosis-inhibited Xenopus laevis. Dev Biol, 142(1), 178–183. DOI: 10.1016/0012-1606(90)90161-b

Serhan, C. N. (2014). Pro-resolving lipid mediators are leads for resolution physiology. Nature, 510(7503), 92–101. DOI: 10.1038/nature13479

Smith, P. B., & Turpen, J. B. (1985). Hemopoietic differentiation potential of cultured lateral plate mesoderm explanted from Rana pipiens embryos at successive developmental stages. Differentiation, 28(3), 244–249. DOI: 10.1111/j.1432-0436.1985.tb00831.x

Smith, S. J., Kotecha, S., Towers, N., Latinkic, B. V., & Mohun, T. J. (2002). XPOX2-peroxidase expression and the XLURP-1 promoter reveal the site of embryonic myeloid cell development in Xenopus. Mech Dev, 117(1-2), 173–186.

Sugiura, T., Tazaki, A., Ueno, N., Watanabe, K., & Mochii, M. (2009). Xenopus Wnt-5a induces an ectopic larval tail at injured site, suggesting a crucial role for noncanonical Wnt signal in tail regeneration. Mech Dev, 126(1-2), 56–67. DOI: 10.1016/j.mod.2008.10.002

Tashiro, S., Sedohara, A., Asashima, M., Izutsu, Y., & Maéno, M. (2006). Characterization of myeloid cells derived from the anterior ventral mesoderm in the Xenopus laevis embryo. Dev Growth Differ, 48(8), 499–512. DOI: 10.1111/j.1440-169X.2006.00885.x

Tseng, A.-S., Carneiro, K., Lemire, J. M., & Levin, M. (2011). HDAC Activity Is Required during Xenopus Tail Regeneration. Plos One, 6(10). DOI: 10.1371/journal.pone.0026382

Tseng, A. S., Beane, W. S., Lemire, J. M., Masi, A., & Levin, M. (2010). Induction of vertebrate regeneration by a transient sodium current. J Neurosci, 30(39), 13192–13200. DOI: 10.1523/JNEUROSCI.3315-10.2010

Tsujioka, H., Kunieda, T., Katou, Y., Shirahige, K., & Kubo, T. (2015). Unique gene expression profile of the proliferating Xenopus tadpole tail blastema cells deciphered by RNA-sequencing analysis. PLoS One, 10(3), e0111655. DOI: 10.1371/journal.pone.0111655

Weller, P. F., Ryeom, S. W., Picard, S. T., Ackerman, S. J., & Dvorak, A. M. (1991). Cytoplasmic lipid bodies of neutrophils: formation induced by cis-unsaturated fatty acids and mediated by protein kinase C. J Cell Biol, 113(1), 137–146. DOI: 10.1083/jcb.113.1.137

