## Supplementary figures and images for "Epigenetic immune-modulation by Histone Deacetylase Activity (HDAC) of tissue and organ regeneration in *Xenopus laevis*"

### Supplemental Figure 1

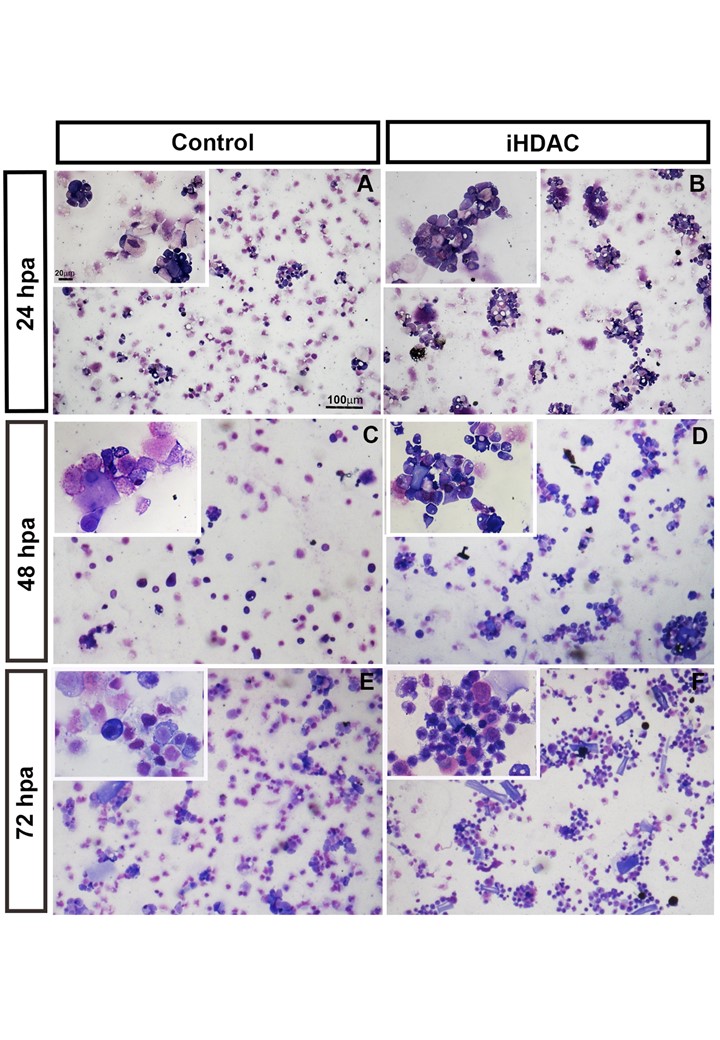

### Supplemental Figure 2

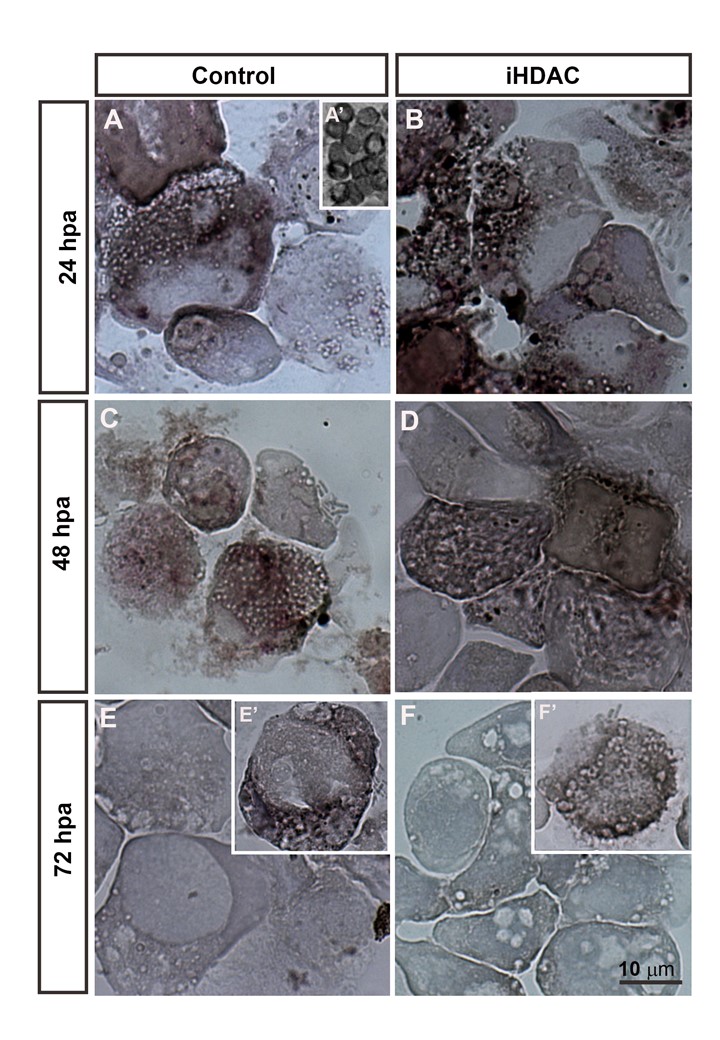

### Supplemental Figure 3

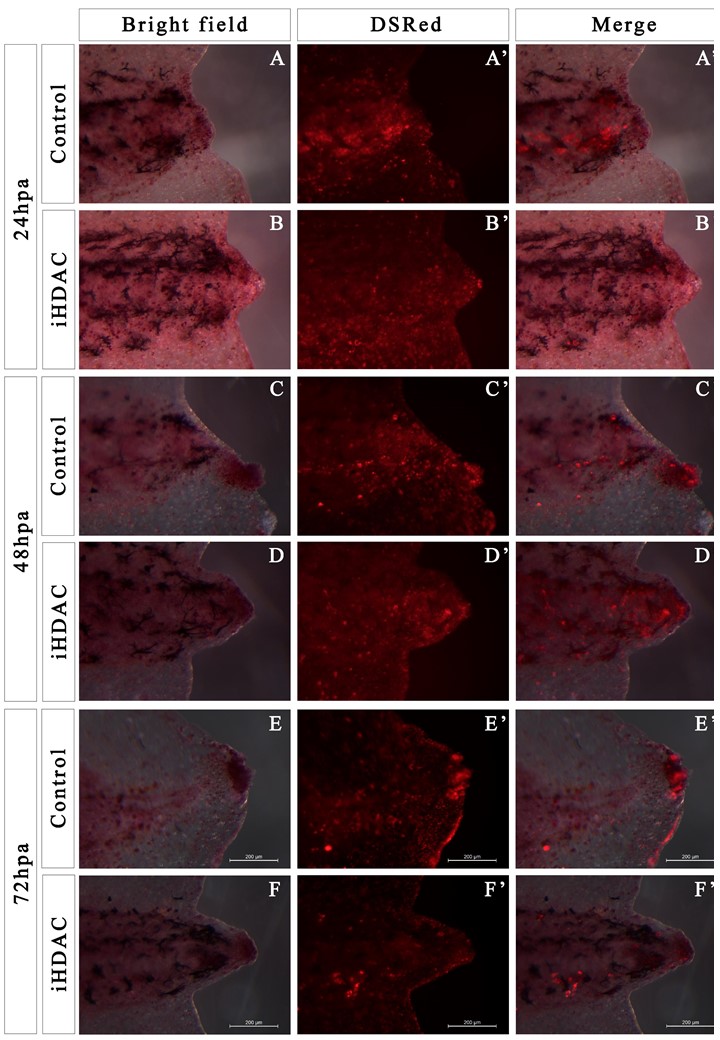

### Supplemental Figure 4

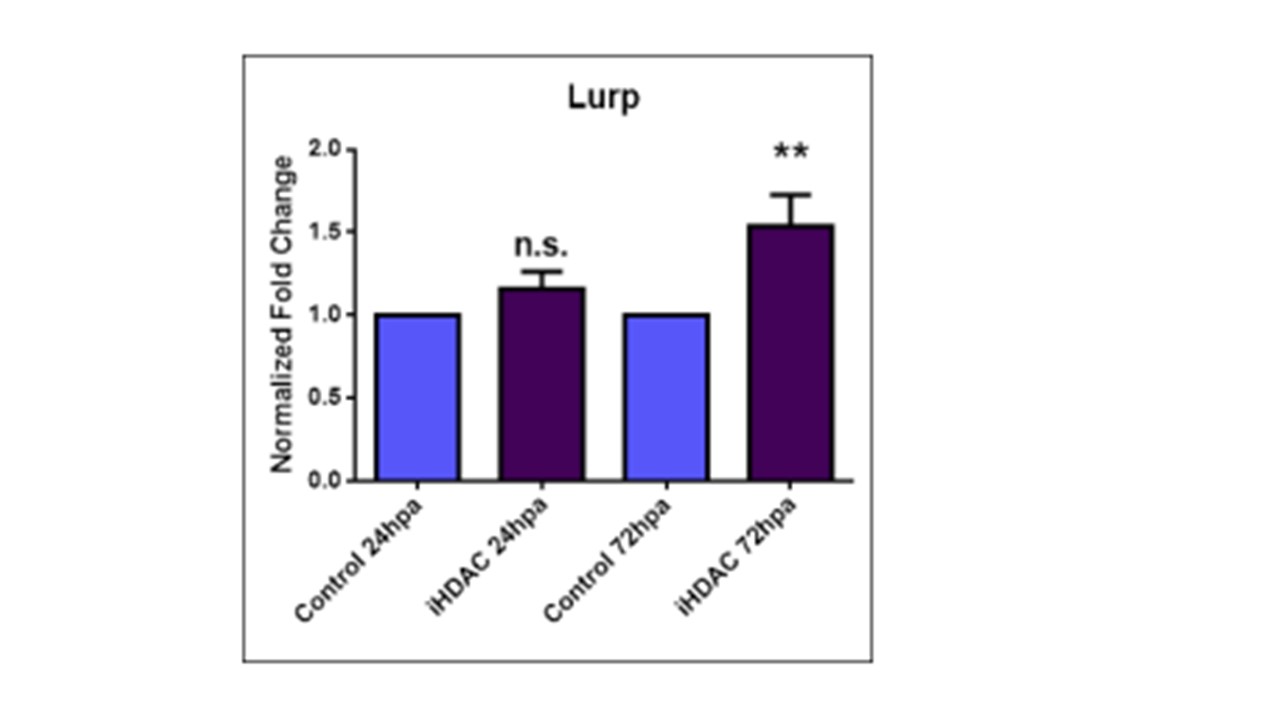
